# High evolutionary and taxonomic diversity of gerbils (*Gerbillus*) in the Horn of Africa

**DOI:** 10.64898/2025.12.09.693121

**Authors:** Danila S. Kostin, Aleksey A. Martynov, Elena V. Cherepanova, Mesele Yihune, Mengistu Wale, Barbora Pavlíčková, Josef Bryja, Leonid A. Lavrenchenko

## Abstract

The genus *Gerbillus* represents an example of the successful Pleistocene radiation among rodents, leading to the emergence of almost 50 currently recognized species inhabiting Asia (from the Middle East to northwest India) and Africa (the Saharo-Sahelian region and eastern Africa). During the last two decades, multiple molecular phylogenetic studies helped to decipher the complex subgeneric structure of this speciose group. However, questions concerning the number of phylogenetic lineages (species) within the genus and their evolutionary relationships remained open, especially because of the lack of genetic material from the Horn of Africa. Here we provide for the first time genetic information on *Microdillus peeli* (being for more than a hundred years a taxon with an unresolved phylogenetic position) and unequivocally show that it is an internal lineage of *Gerbillus*. Together with one additional newly characterized lineage of apparently subgenus rank (proposed here as subgenus “*Pusillus*”), endemic to Somali-Masai region in eastern Africa, these findings document that the Horn of Africa is the center of evolutionary diversity of gerbils in general, and of the genus *Gerbillus* in particular.

## Introduction

Leveraging of molecular genetic data has opened wide opportunities to order our knowledge concerning biological diversity. However, even among mammals, there are still many groups with unresolved taxonomic questions, persisting as a result of either an absence of material suitable for molecular analysis or the existence of cryptic lineages of unclear taxonomic status. Here, we focus on the members of subtribe Gerbillina (subfamily Gerbillinae, tribe Gerbillini; review in Pavlinov, 2008), a clade of African–southern Asian gerbils, which currently includes three genera (Pavlinov, 2008; Wilson et al., 2017): *Gerbillus*, *Sekeetamys,* and *Microdillus*. While the last two appear to be monotypic, the former one possesses a complex structure and high species diversity (Ndiaye et al., 2016).

Representatives of the genus *Gerbillus* span a wide distribution across the African-Palearctic belt of arid zones ranging from western Africa to north-western India (Ndiaye et al., 2016). However, most studies on the evolutionary systematics of this group involved a limited number of species, mainly from Western and Northern Africa (Alhajeri, 2018; Cui et al., 2025). Members of the genus *Gerbillus* inhabiting the so-called Horn of Africa (current territories of Ethiopia, Somalia, Djibouti, and Kenya) were not included in most complete recent phylogenetic studies of gerbils (Ndiyae et al., 2016; Piwczyński et al., 2023), although it concerns a biogeographically important region for arid-adapted organisms (Bryja et al., 2022), considered as a cradle for the whole subfamily Gerbillinae (Kostin et al., 2022). Four well-defined genetic clades within the genus *Gerbillus* were revealed by the most comprehensive (in terms of number of analyzed specimens, i.e. 89 individuals from 71 localities) phylogenetic analysis of the genus: *Gerbillus*, *Dipodillus*, *Monodia*, and *Hendecapleura* (Ndiyae et al., 2016). Taking into account their relatively recent origin (between 5.1-3.7 Mya; Ndiaye et al., 2016), the authors referred to these lineages at the subgeneric, rather than the generic level. Being based on a minimal set of molecular markers (one mitochondrial and one nuclear gene), the work of Ndiyae et al. (2016) still produced a resolved topology, which, in general, was confirmed later by using a set of 2,650 ddRAD-seq loci (Piwczyński et al., 2023). Thus, the most basal position occupies *Hendecapleura* (comprising *G. nanus*, *G. amoenus,* and *G. henleyi*). Further divisions into *Gerbillus*, *Dipodillus*, and *Monodia* happened almost simultaneously, and alternative topologies for this triad were revealed in different phylogenetic reconstructions, showing either a basal position of *Dipodillus* (Ndiaye et al., 2016) or *Monodia* (Piwczyński et al., 2023).

Position of *Sekeetamys*, the monotypic genus with the only species, *S. calurus* (Thomas, 1892), has been uncertain for a long time due to the intermediate nature of its morphological traits. It has been considered either as the most advanced member of Gerbillina or the most primitive member of Rhombomyina (Pavlinov, 2008). Later, available molecular data supported the first hypothesis placing *Sekeetamys* as a sister group to the members of the *Gerbillus* genus (Chevret & Dobigny, 2005; Ndiyae et al., 2016; Kostin et al., 2022).

The last lineage within the Gerbillina subtribe, whose precise phylogenetic position remains unknown, is *Microdillus peeli* (de Winton, 1898) – a small gerbil, described by William Edward de Winton based on a single individual, collected by Mr. C.V.A. Peel in Somaliland. Already in the species description, de Winton noted special morphological features distinguishing this species: “*The skull is peculiarly square and short and unlike any other Gerbil I know*”. Later, Pavlinov (2008), in his review on phylogeny and classification of Gerbillinae, mentioned this species as being close to the members of the genus *Gerbillus*, but at the same time made the reservation that this species’ morphological evolution apparently went into another direction compared to other typical *Gerbillus*, which supports the generic rank of the taxon (Pavlinov, 2008). To date, molecular data for *M. peeli* have been absent due to the scarcity of the species in both field and museum collections. This resembles the fate of another monotypic gerbil genus – *Ammodillus*, with a single species *A. imbellis* (de Winton, 1898) – described in the same paper with the *M. peeli*, but only relatively recently placed on the phylogenetic tree using molecular data (Bryja et al., 2022; Kostin et al., 2022). Here, after more than 125 years since *M. peeli* discovery and formal description, we provide for the first time the data on its karyotype and mitochondrial and nuclear markers and place it onto the molecular phylogenetic tree of gerbils. Furthermore, we provide new data and characterize an additional lineage of *Gerbillus* from the Horn of Africa, previously proposed as a putative separate subgenus on the basis of mitochondrial DNA phylogeny (Bryja et al., 2022).

## Material and methods

### Sampling

The study utilized specimens collected during Joint Ethio-Russian Biological Expedition and several Czech research projects (see more details on collectors and localities in Supplementary Table S1). Distribution data for the collected specimens are presented in Fig. 1. Three *M. peeli* individuals were collected in 2022 in eastern Ethiopia, and one in Somaliland (collected by Tomáš Mazuch in 2018). Other *Gerbillus* individuals used in the study were collected in southern and eastern Ethiopia, central Kenya, and northern Tanzania (Fig. 1, Supplementary Table S1). Tissue samples were preserved in 96% ethanol and stored at 4°C in the collections of the A. N. Severtsov Institute of Ecology and Evolution (Russian Academy of Sciences, Moscow, Russia) and the Institute of Vertebrate Biology (Czech Academy of Sciences, Brno, Czech Republic). Voucher specimens, including skulls and dry-prepared skins, were deposited in the Zoological Museum of Moscow State University (ZMMU, Moscow, Russia) and the Institute of Vertebrate Biology (Brno, Czech Republic).

**Fig. 1.**
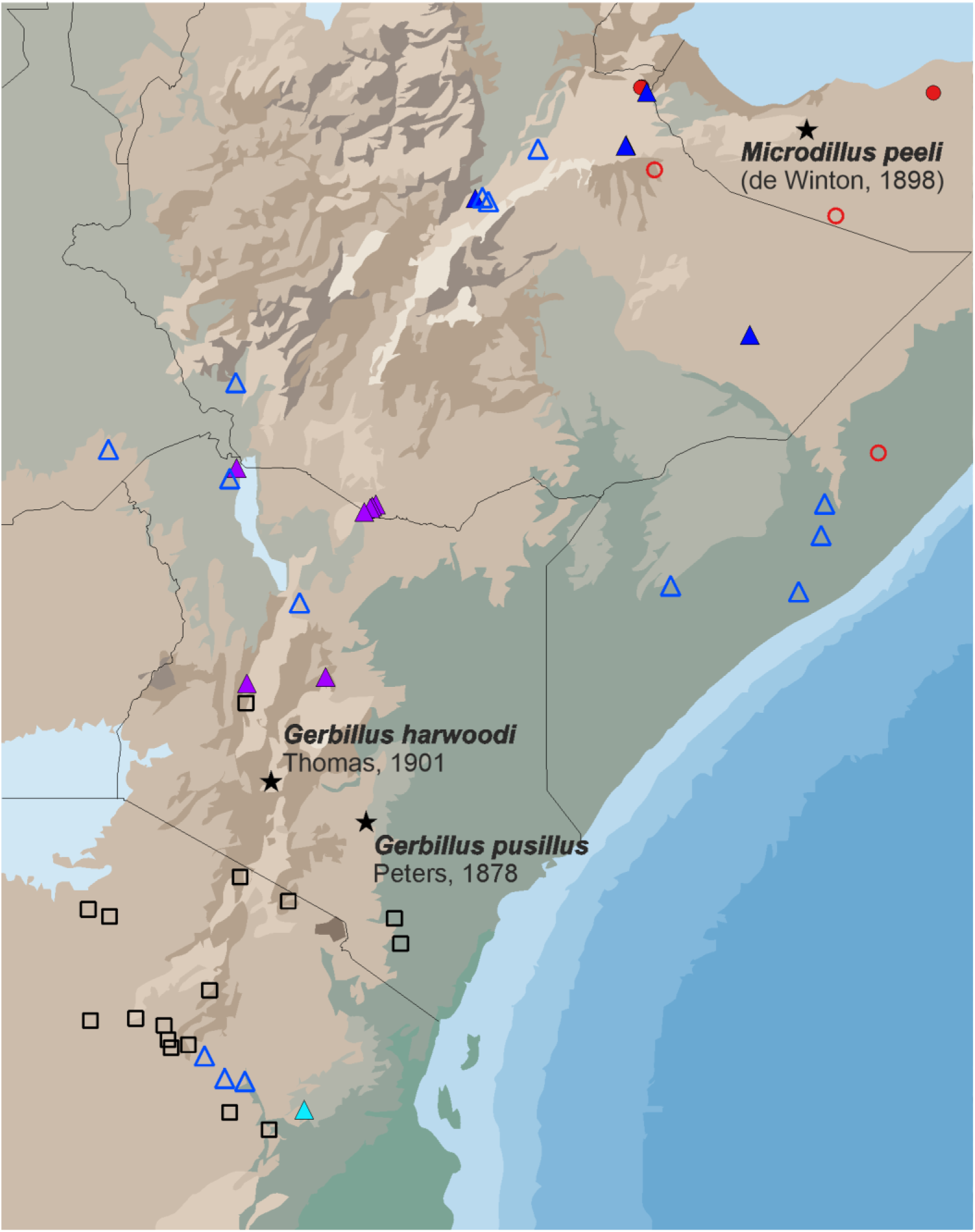
Distribution map of *M. peeli* (red) and *G. pusillus* (blue – ‘Ethiopian’, purple – ‘Ethio-Kenyan’, and turquoise - ‘Tanzanian’ lineages). Filled figures denote genotyped individuals, empty figures – ungenotyped records following Monajem et al. (2015). Black empty squares denote records reported as *G. harwoodi*. Black stars denote the type localities of the mentioned taxa.

### DNA Extraction and Sequencing

DNA was extracted using a commercial kit following the manufacturer’s protocol. Mitochondrial cytochrome *b* (*cytb*) and four nuclear markers—tartrate-resistant acid phosphatase (*Acp5*), growth hormone receptor (*GHR*), interphotoreceptor retinoid-binding protein (*IRBP*), and recombination-activating gene 1 (*RAG1*)—were amplified and sequenced with the following set of primers (5’-3’): *cytb:* L14723 [ACCAATGACATGAAAAATCATCGTT] and H15915 [TCTCCATTTCTGGTTTACAAGAC]*; Acp5:* 120FWD [AATGCCCCATTCCACACAGC] and 564REV [GCAGAGACGTTGCCAAGGTG] (DeBry and Seshadri, 2001); *GHR:* GHR1F [GGRAARTTRGAGGAGGTGAACACMATCTT] and GHR2R [GATTTTGTTCAGTTGGTCTGTGCTCAC] (Adkins et al., 2001); *IRBP:* IRBP217F [ATGGCCAAGGTCCTCTTGGATAACTACTGCTT] and IRBP1531R [CGCAGGTCCATGATGAGGTGCTCCGTGTCCTG] (Stanhope et al., 1992); *RAG1:* Rag1_F50 [GGGATGGGGGATGTGAGTGAGAAG] and Rag1_R2864 [TTTCAGGGACATGTGCCAAGGTTTT] (Zemlemerova et al., 2021). The *cytb* gene was amplified using the protocol described by Lecompte et al. (2002). A touchdown PCR program was employed for nuclear markers, with the following conditions: initial denaturation at 94°C for 3 min; 40 cycles of 94°C for 40 s, 67–57°C (10 cycles starting at 67°C, decreasing by 1°C per cycle, followed by 30 cycles at 57°C) for 45 s, and 72°C for 1 min. PCR products were sequenced using the Sanger method. Reference sequences for *cytb* and nuclear markers from members of each subgenus of *Gerbillus* were retrieved from GenBank. The list of all sequences employed in this work, including the GenBank accession numbers of newly obtained sequences, is provided in Supplementary Table S2. The species and subgeneric names used in this work were taken following the most recent phylogenetic papers of Ndiaye et al. (2016) and Piwczynski et al. (2023).

### Phylogenetic Reconstructions

Nucleotide sequences were edited and aligned using BioEdit v.7.0.5.3 (Hall 1999). Pairwise genetic *p*-distances and between-group mean p-distances were calculated from the *cytb* alignment in MEGA X v.10.2 (Kumar et al. 2018). Phylogenetic reconstructions were performed separately for mitochondrial and nuclear data using a maximum likelihood and Bayesian approaches. A maximum likelihood (ML) tree was inferred using IQ-TREE v.2.3.6 (Nguyen et al. 2015), with partitioning strategies and substitution models selected via ModelFinder (Kalyaanamoorthy et al. 2017). Clade support was assessed using 5000 bootstrap replicates. Bayesian inference (BI) was performed in MrBayes v.3.2.6 (Ronquist et al. 2012) with three heated chains and one cold chain, starting from random trees and consisting of 5,000,000 generations. The best-fit partitioning scheme and substitution models were determined using PartitionFinder v.2.1 (Lanfear et al. 2017).

### Species Tree and Molecular Dating

A species tree based on four nuclear markers was reconstructed using the Bayesian framework implemented in *BEAST 3 (Douglas et al., 2022), an extension of BEAST 2 (Bouckaert et al., 2019). Gene alignments were processed in BEAUti 1.8.2, with separate and unlinked substitution, clock, and tree models assigned to each marker. The partitioning strategy and substitution models were selected using ModelFinder. The tree was modeled under a birth-death process with lognormal relaxed clocks and three fossil-based calibration points applied as follows: a) divergence between tribes Gerbillurini (*Desmodillus*, *Gerbilliscus*, and *Tatera*) and Gerbillini (*Gerbillus*, *Microdillus*, *Sekeetamys*, *Taterillus, Rhombomys*, *Meriones*, *Psammomys*, and *Brachiones*), based on *Abudhabia pakistanensis* (8.6 Mya), see Kostin et al., 2022 for notes on this calibration point; b) the earliest records of *Gerbilliscus* from Lemudong’o, Kenya (Manthi, 2007), corresponding to the divergence between *Desmodillus* and *Gerbilliscus* (6.1 Mya); c) divergence of *Gerbillus* from other Gerbillinae lineages, based on the oldest representative of this genus from the early Pliocene of Kenya (4 Mya). Here, we follow Ndiyae et al. (2016), interpreting this point as a MRCA of *Gerbillus*-*Sekeetamys*. Calibration densities followed a lognormal distribution for all three points (mean = 1.6, SD = 1) with offset equal to the age of particular fossil. Two independent runs of 100 million generations were performed, sampling every 10,000 generations. The first 15% of samples were discarded as burn-in, and convergence was assessed using Tracer 1.6 (Rambaut et al. 2014). Results were combined using LogCombiner (v2.7.7), and a maximum clade credibility tree was generated with TreeAnnotator (v2.7.7) included in the BEAST 2 software (Bouckaert et al., 2019).

### Karyotyping

Samples of metaphase plates have been obtained from the bone marrow of three individuals of *G. pusillus*, and, for the first time, from the three Ethiopian specimens of *M. peeli* using a classical procedure (Ford & Hamerton, 1956). Giemsa staining was performed in 4% solution.

### Geometric Morphometrics

Skull shape and size variations were analyzed using a two-dimensional geometric morphometric approach. To maximize the coverage of the skull shape variability among the members of the *Gerbillus* genus, we analyzed 75 skulls from 10 species (see details in Supplementary Table S1). Only complete skulls of adult specimens were included. Dorsal, ventral, and lateral skull images were captured with a Pentax K-3 digital camera equipped with a macro lens. Skull shape was characterized using a total of 94 anatomical landmarks (Supplementary Fig. S1), digitized in TpsDig 2.12 (Rohlf, 2008). Landmark configurations for each view were superimposed via generalized Procrustes analysis (Rohlf & Slice, 1990) to eliminate non-shape variation (rotation, scale, and position). Procrustes shape coordinates and centroid size were separately extracted for further analysis, and matrices of partial warp scores of shape were computed in TPSrelw (Rohlf, 2007). Including centroid size values as a column to the shape scores matrix allows performing further analysis either separately on shape or size, or combining shape and size within a joint framework. Principal component analysis (PCA) was performed on shape matrices for each skull view (dorsal, lateral, ventral) and on a combined dataset. Analyses were conducted in PAST 4.11 (Hammer et al., 2001), examining both shape with centroid size and shape alone. Multivariate linear regression was performed to assess the relationship between the skull’s log centroid size and PCs of shape.

## Results

### Phylogenetic analyses

Both mitochondrial and nuclear trees (Fig. 2) provide similar topologies and possess six highly supported lineages of comparable degree of evolutionary divergence in the genus *Gerbillus*. They include four previously delimited subgenera: *Dipodillus*, *Gerbillus*, *Monodia*, and *Hendecapleura* (e.g., Ndiaye et al., 2016), as well as the so-called ‘*Pusillus*’ group, comprising individuals from Kenya, Tanzania, and southern and eastern Ethiopia (in agreement with our previous work; Bryja et al., 2022; Kostin et al., 2022). For simplicity, we will call this clade as “*Pusillus*” hereafter; see below for more taxonomic details). Most surprisingly, all newly produced sequences of the genus *Microdillus* form a similarly distinct sixth branch within *Gerbillus*, sister to the subgenus *Hendecapleura* in all analyses. The mean *p*-distances at *cytb* marker between the six major lineages (i.e., proposed subgenera) are equal to 12-17% (Table 1).

**Fig. 2.**
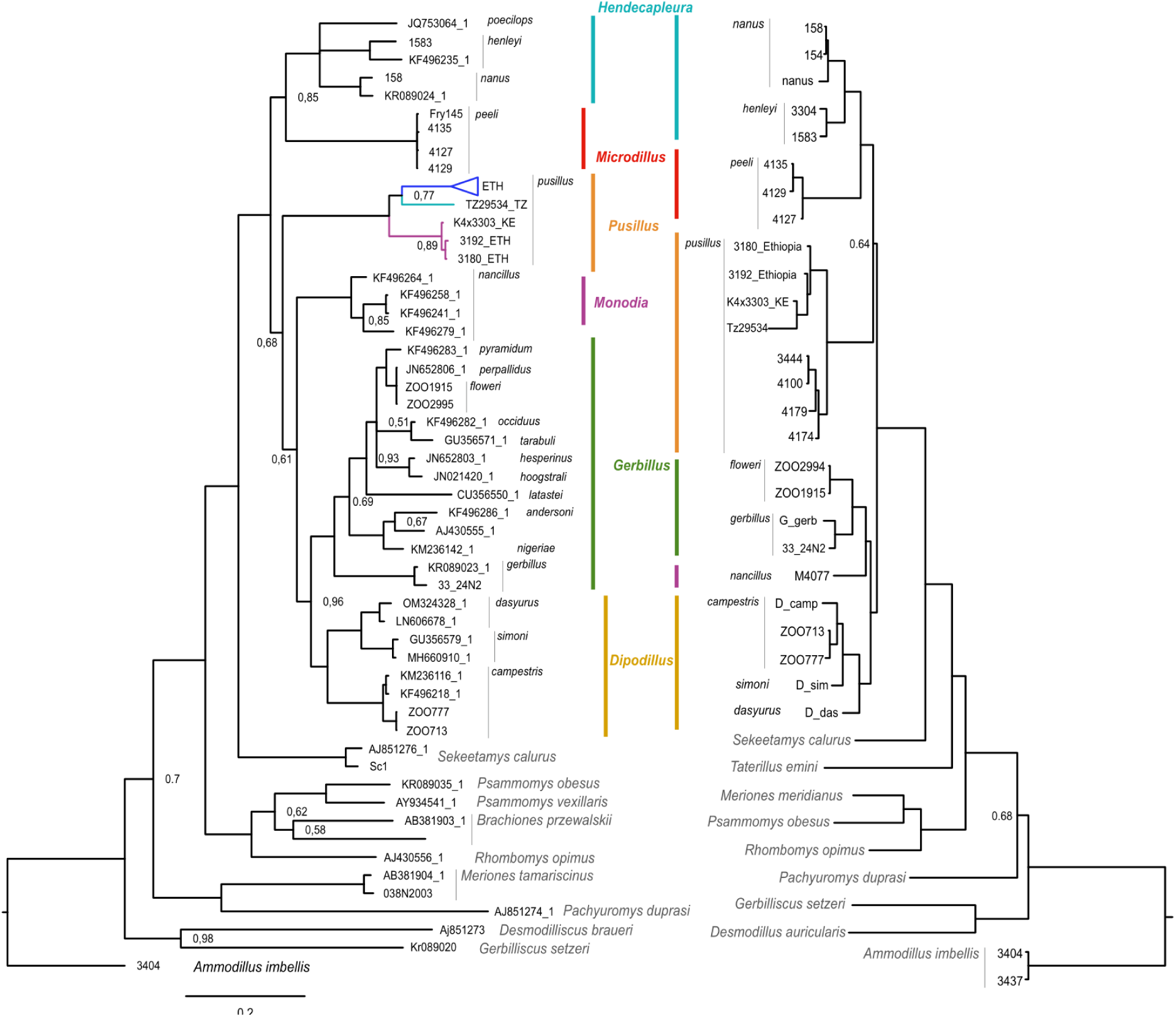
Bayesian phylogenetic trees based on *сytb* gene sequences (left) and four concatenated nuclear markers (right). Posterior supports equal to 1.0 are not shown. Colored dotted lines define six sub-generic lineages within the *Gerbillus* genus. Separate branches within *G. pusillus* on mtDNA tree colored according to Fig. 1.

**Table 1.** Mean genetic *p*-distances at *cytb* among genetic lineages within *Gerbillus* genus (below diagonal). Standard errors of inter-lineage distances calculated by 1 000 bootstraps are above the diagonal.

|  | *Pusillus* | *Hendecapleura* | *Microdillus* | *Monodia* | *Gerbillus* | *Dipodillus* |
| --- | --- | --- | --- | --- | --- | --- |
| *Pusillus* |  | 0,009 | 0,012 | 0,010 | 0,010 | 0,010 |
| *Hendecapleura* | 0,153 |  | 0,010 | 0,009 | 0,008 | 0,008 |
| *Microdillus* | 0,177 | 0,148 |  | 0,010 | 0,009 | 0,009 |
| *Monodia* | 0,153 | 0,143 | 0,147 |  | 0,008 | 0,008 |
| *Gerbillus* | 0,157 | 0,140 | 0,148 | 0,132 |  | 0,007 |
| *Dipodillus* | 0,164 | 0,137 | 0,146 | 0,126 | 0,123 |  |

Results of the divergence time estimation are visualized on the dated species tree (Fig. 3). The crown age of the Gerbillina subtribe represented by the split of lineages leading to *Sekeetamys* and *Gerbillus* genera is estimated around 4.6 Mya (4.1-5.3). Subsequent splits happened almost simultaneously in the lower Pleistocene and led to the emergence of six major lineages within the *Gerbillus* genus.

**Fig. 3.**
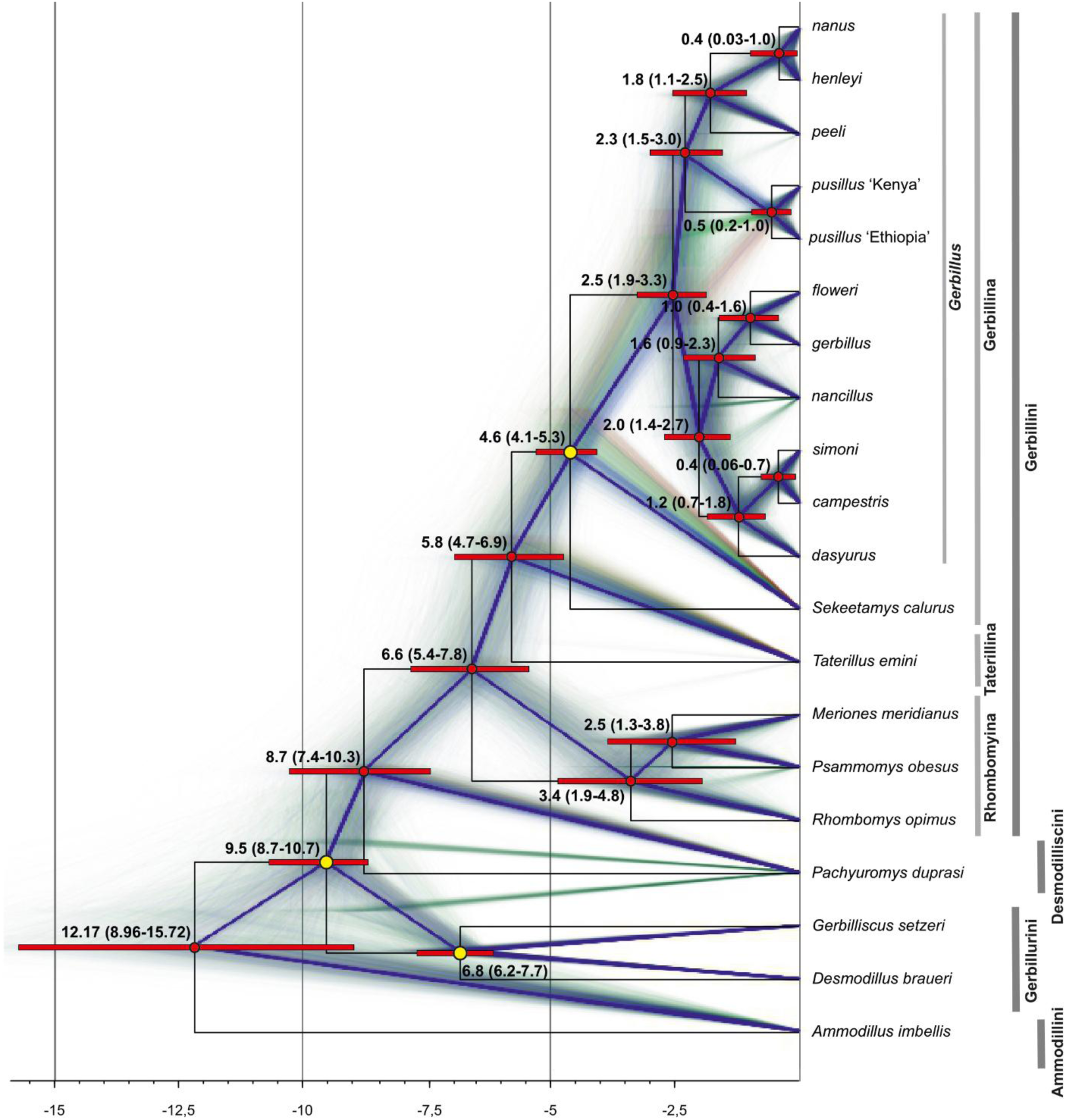
Dated species tree (based on four nuclear markers) with a visualization of 1 500 sampled trees in the background. Values on the nodes denote estimated divergence age and 95% confidence intervals. Yellow circles denote the position of the fossil-based calibration points. Vertical bars show the taxonomic division within the subfamily.

### Chromosomal analysis

The karyotype of female individual of *M. peeli* shows a diploid number with 2n = 44, NF = 54, and comprises one pair of large metacentrics, one pair of large submetacentrics, three pairs of medium metaсentrics and submetacentrics, gradually diminishing in size, and 17 pairs of acrocentrics, gradually decreasing from medium to small (Fig. 4). The sex chromosomes were not determined. The karyotype, obtained for a *G. pusillus* female individual from Kebri-Dahar (eastern Ethiopia), is characterized by 2n = 34, NF = 54 (8 metacentric, one submetacentric, seven acrocentric autosomal pairs; X chromosome represented by a large submetacentric), which is apparently identical to one described for *G. pusillus* from Somalia (Capanna & Merani, 1981). The minor differences were related to the sex chromosomes. In Cappana & Merani’s work, X chromosomes are large acrocentrics. However, taking into account full compliance in the number and shape of the autosomes, as well as moderate quality of the obtained plates, it can be assumed that displayed differences represent a random artifact related more to the different degrees of chromosome condensation, rather than to a real difference in the shape of the sex chromosomes.

**Fig. 4.**
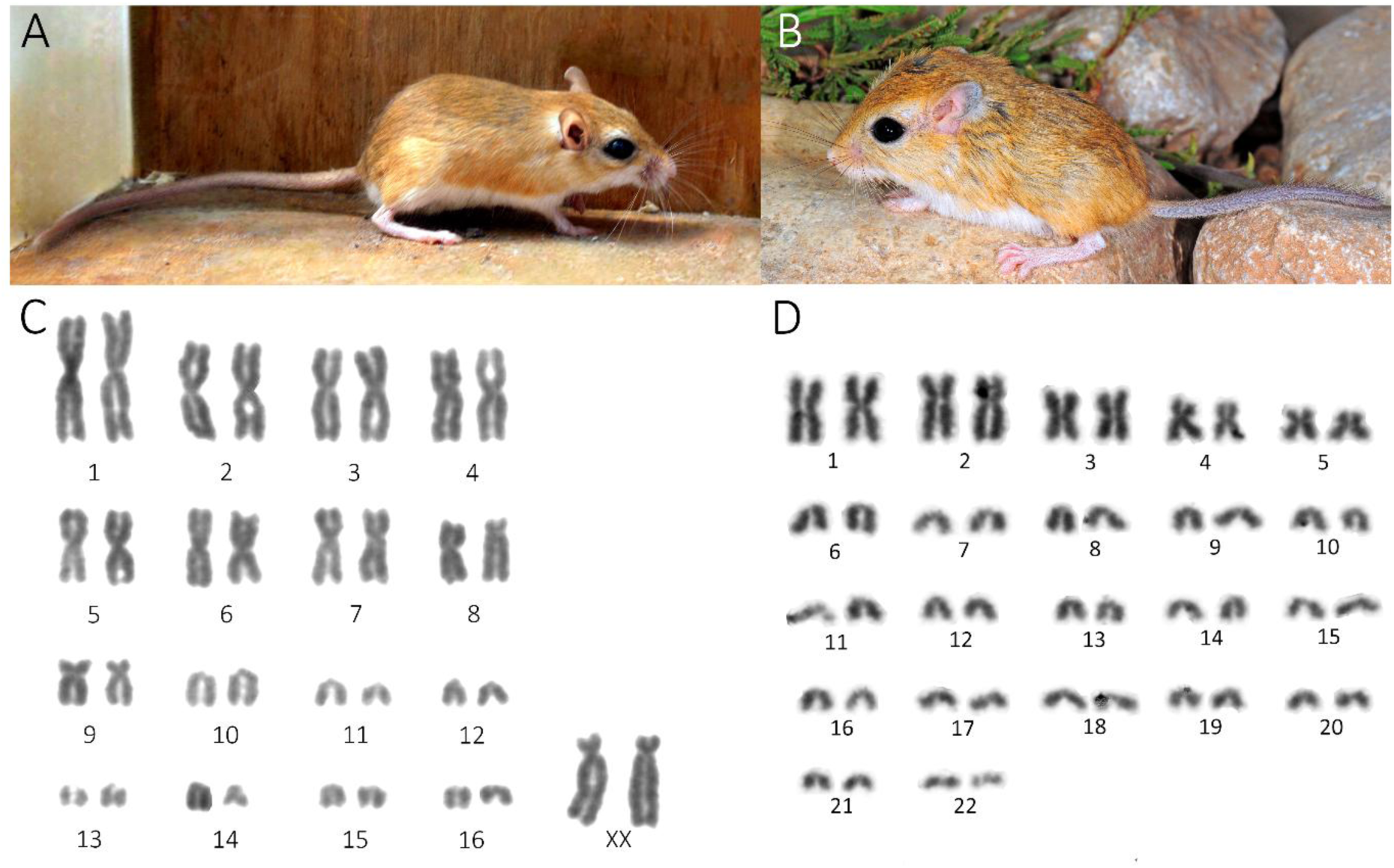
External appearance (photos by Alexey Martynov and Tomáš Mazuch) and karyotypes for *G. pusillus* (A, C) and *M. peeli* (B, D).

### Morphology analysis

The species analyzed are well differentiated in the morphospace based on their skull shape (Fig. 5), however the clustering revealed does not correspond to their phylogenetic relationships with the division into six lineages. Instead of this, the order in which species are arranged along the first two principal components is highly correlated (r=0.74, p<0.001 for PC1 and r=0.39, p=0.01 for PC2) with the centroid size of skull, independently obtained within the separate analysis (Fig. 5 B-C).

**Fig. 5.**
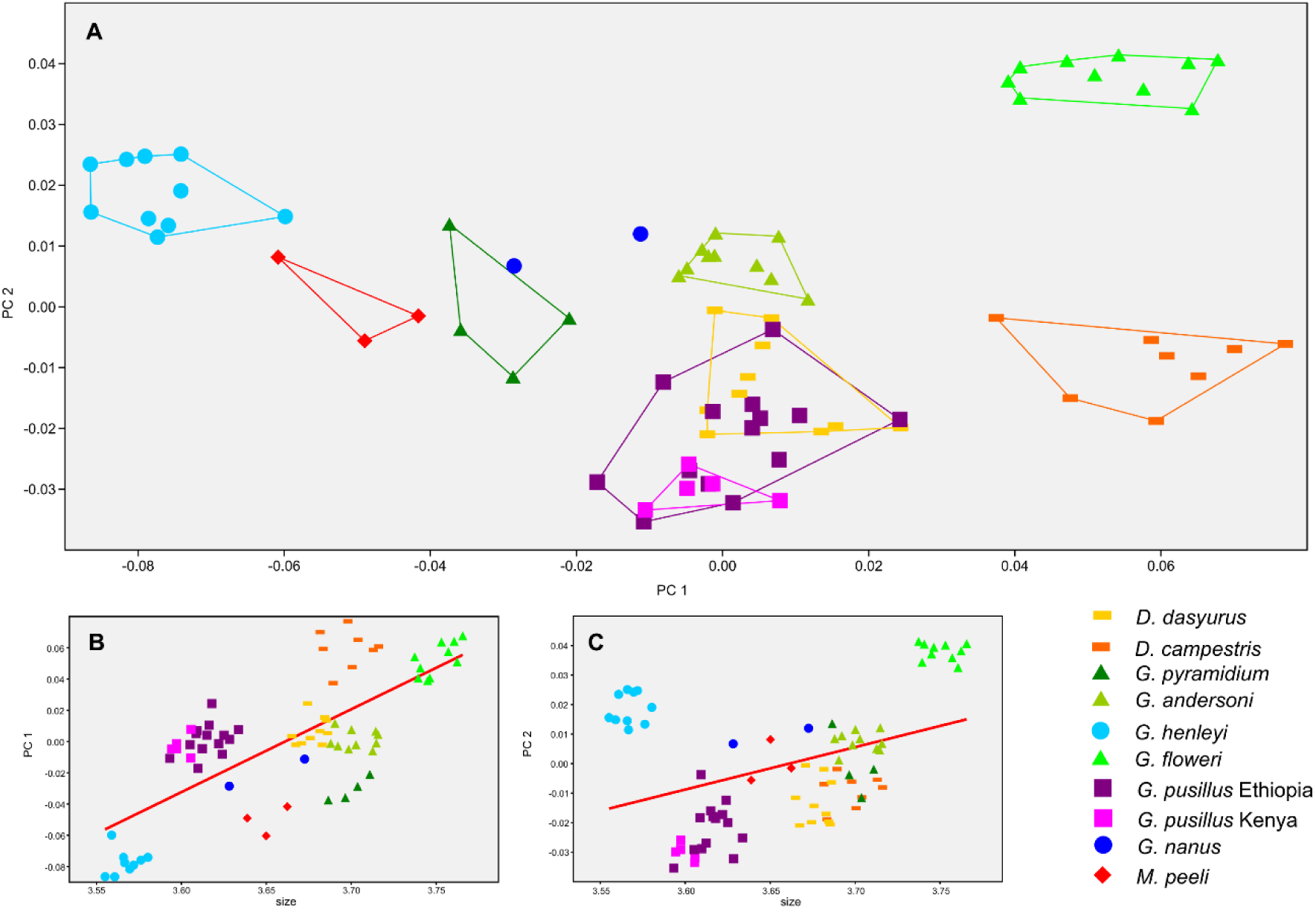
A. Skull shape scatter plot in the space of the first two principal components. The order in which species are arranged along the first two principal components has found high correlation with the obtained centroid size of skull for PC1 (B) and moderate correlation for PC2 (C).

## Discussion

*M. peeli* and *G. pusillus* are internal deeply divergent lineages within the *Gerbillus* genus Analysis of both mitochondrial and nuclear data confidently placed *M. peeli* as an inner, deeply divergent lineage within the *Gerbillus* genus. Along with this, *M. peeli* possesses unique features (strikingly different diploid number and a whole range of morphological, cranial, and dental traits), distinguishing it not only from its sister clade *Hendecapleura* (represented by *G. henleyi*, *G. nanus*, and *G. amoenus*) but also from other members of the *Gerbillus* genus (Table 2). However, based on our morphology analysis utilizing a geometric morphometric approach, *M. peeli* falls within the skull shape variation of the *Gerbillus* genus (Fig. 5).

**Table 2.**
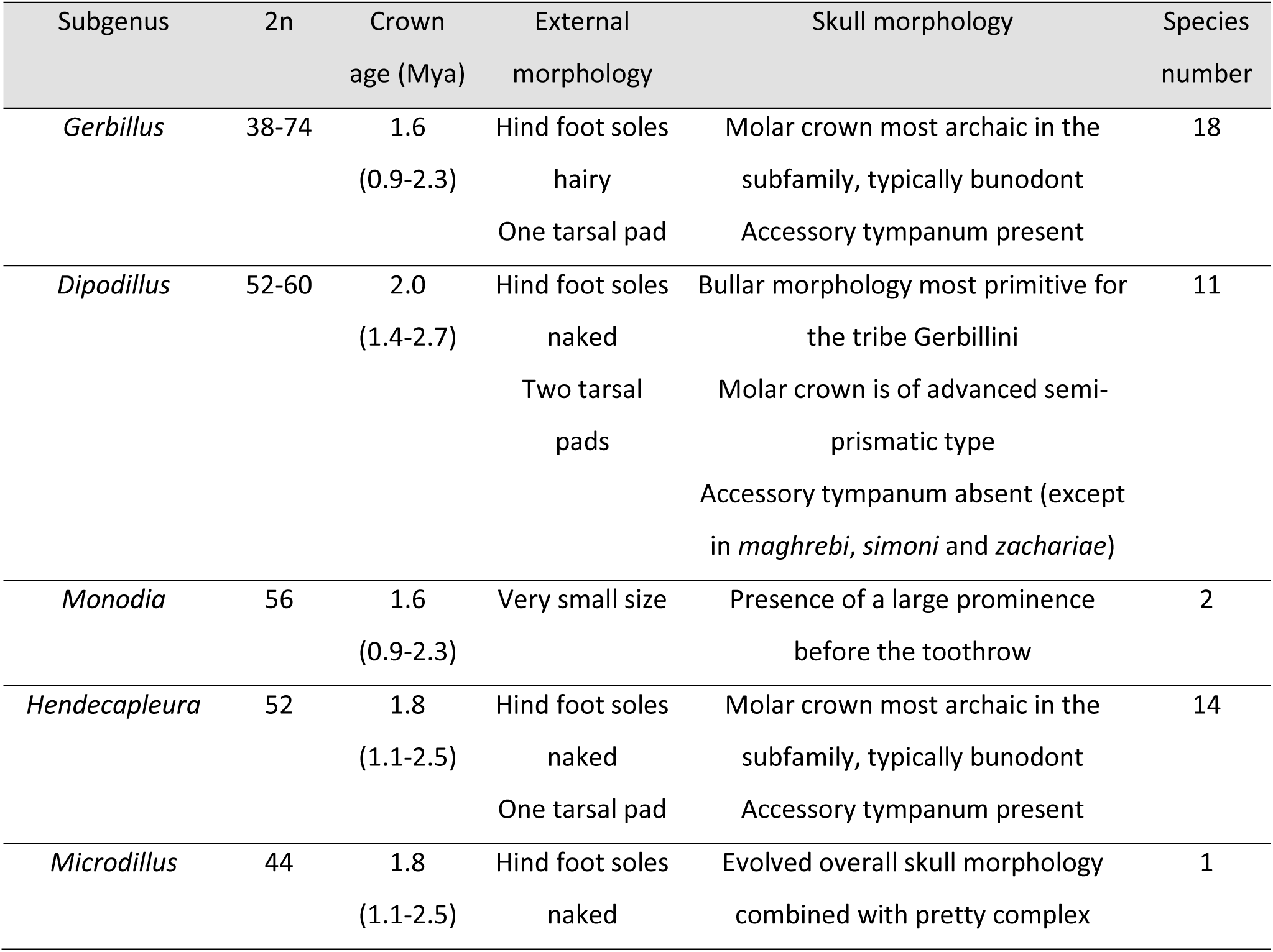
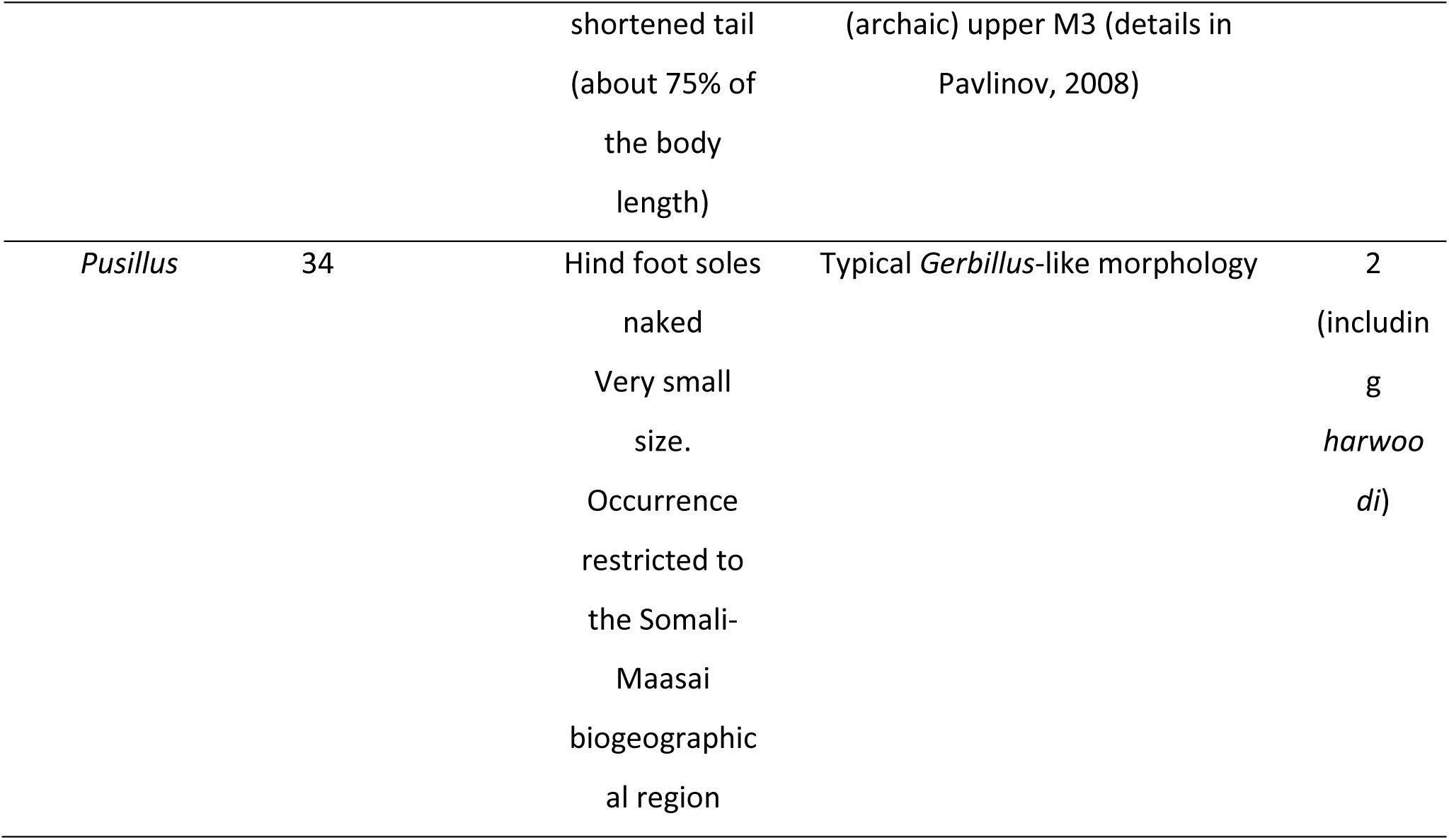
Comparative characteristics of the six subgenera within defined within the *Gerbillus* genus. Chromosome diploid number (2n) is given following Granjon, 2013. External and Skull morphology is given following Ndiyae et al., 2016 (with additions). Species number is given following Mammal Diversity Database (2025). All data modified with our changes regarding taxa under study.

The other finding of our phylogenetic analysis is that small gerbils from Eastern Africa (Kenya, Tanzania, and southern and eastern Ethiopia) also represent another independent offshoot within the *Gerbillus* genus stem (see also Bryja et al, 2022). We attributed them to *G. pusillus* based on chromosomal analysis of individuals from eastern Ethiopia, revealing a karyotype with the smallest diploid number within the genus (2n =34, Fig. 4), identical to one, previously reported for this species (Capanna & Merani, 1981).

### Six subgeneric clades within the *Gerbillus* genus

The precise dates of the crown *Gerbillus* genus vary in different works (5.1 Mya – Ndiyae et al., 2016; 2.5 Mya – Kostin et al., 2022; 4.66 Mya - Piwczyński et al., 2023; 2.5 Mya – this study). We are prone to consider that some of these estimations are significantly overestimated due to the use of mitochondrial DNA sequences (or secondary calibration points, based on the mtDNA sequences analysis), that usually makes nodes “older” due to a significantly higher mutation rate in the mitochondrial genome. The other issue of paramount importance for accurate dating divergence is a using of correct calibration points. Despite the presence of several dated fossils within the group, their correct placement on the Gerbillinae subfamily phylogenetic tree still remains an open question and vary in different works. Thus, fossil of *Abudhabia pakistanensis*, known from the Late Miocene deposits, apparently being a Gebillurini stem taxon, treated either as crown for the Gerbillurini (Aghová et al., 2017) or as TMRCA of Gerbillini – Gerbillurini split (Kostin et al., 2022).

Following our data, inclusion of *Microdillus* and “*Pusillus*” into the *Gerbillus* phylogenetic tree revealed the first split of the ancestral stem into three lineages, namely ‘*Microdillus* + *Hendecapleura’*; ‘*Monodia* + *Dipodillus* + *Gerbillus’*; and ‘*Pusillus’*, and practically simultaneous subsequent divisions, leading to emergence of the six lineages. Being initially described as separate genera, *Monodia*, *Dipodillus*, and *Hendecapleura* has been moved as inner lineages in *Gerbillus* genus based on the results of molecular analysis and to date considered as subgenera in order to emphasize complex structure of the *Gerbillus* genus (Ndiaye et al., 2016). Following our results, given the similar genetic divergence level of *Microdillus* and “*Pusillus*”, these two lineages deserve the same rank (i.e. to downgrade *Microdillus* to subgenus and describe new subgenus “*Pusillus*”), as other subgenera, previously defined within the *Gerbillus* genus, namely *Gerbillus*, *Dipodillus*, *Monodia*, and *Hendecapleura*. The formal taxonomy of subgenera within *Gerbillus* presents difficulties, since their original species composition (adopted in their description as independent genera) is poorly consistent with the data of modern molecular phylogeny. For instance, both species of proposed subgenus “*Pusillus*” (*G. pusillus* and *G. harwoodi*) are traditionally classified within the two other subgenera - *Hendecapleura* and *Dipodillus*, respectively (Pavlinov et al., 1990; Mammal Diversity Database, 2025). At the same time, it is worth to note, that despite having undoubted evidences favoring separate position of the “*Pusillus*” lineage, within the context of this study we refrain from its formal description as subgenus, due to the difficulties in identifying reliable morphological criteria for delimiting all proposed subgenera, and lack of genetic information preventing association of already described taxa with a candidate type species which would propose for subgenus “*Pusillus*” (see below).

In contrast to molecular data, other criteria as external, craniological and chromosomal data, while widely used in gerbils’ systematics, сan hardly be clearly associated with one specific group (Table 2; see also Supplementary Fig. S2 with the rescaled images of dorsal and lateral skull projections for the different members of the *Gerbillus* genus). Despite that some members of the genus indeed possess remarkable cranial and dental traits (Pavlinov, 2008; Ndiaye et al., 2016), we failed to discriminate members of different subgenera using the geometric morphometry approach (Fig. 5A), which is similar to the previous attempts (Ndiaye et al., 2012; Alhajeri et al., 2018). However, we revealed a pattern of shape differences (i.e. slightly wider cranium, relatively shortened rostrum) among species being correlated with an increase in the centroid size of skull instead of the species’ taxonomic relationships (Fig. 5 B-C).

Despite strong support for all six revealed lineages on the phylogenetic tree, their mutual relationships in some nodes have low support and produce alternative topologies between mitochondrial and nuclear datasets, or while using different methods of phylogenetic tree inference. The first such case is the relationships among *Gerbillus*, *Dipodillus*, and *Monodia*, where the basal group is represented by the *Monodia* (mtDNA) or *Dipodillus* (nuclear data). Exactly the same pattern has been shown in the work of Ndiaye et al. (2016), which is not surprising given the use of overlapping data sets. In the recent work that implemented genomic SNPs (Piwczyński et al., 2023), basal group within this ternary is *Monodia*, however, again with low bootstrap support, also implying an alternative topology. Thus, one can assume nearly simultaneous divergence of these three lineages from the common ancestor. The second case is the poorly resolved position of “*Pusillus*” lineage, clustering closely with the ‘*Gerbillus* + *Dipodillus* + *Monodia*’ clade (mtDNA and ML nuclear tree) or with the ‘*Microdillus* + *Hendecapleura*’ clade (BI nuclear tree). One can assume that the application of the high-resolution genomic data will solve this issue and reveal more precisely the order of splits in the basal radiation of the genus *Gerbillus*.

### Cryptic diversity within the “*Pusillus*” lineage

Species diversity of the East African pygmy gerbils, characterized by the bare hind foot soles but normal tympanic bulla (sensu Roche, 1975), is still a matter of question due to the lack of molecular data from available type specimens. Based on the results of morphological and colorimetric analyses of specimens from Ethiopia and Kenya attributed to *G. pusillus*, *G. ruberrimus*, *G. diminutus*, and *G. percivali*, Roche (1975) proposed synonymizing all of them under the name *G. pusillus*. However, he still preferred to keep a separate species status of *G. harwoodi*, referring to Thomas’ description (1901) and several distinguishing traits, including overall appearance, craniological, and fur colour characters. At the same time, he noted that some individuals possessed intermediate features, which makes their precise species identification “often problematic, even random”. Our phylogenetic analysis revealed three mitochondrial lineages (two of them are well supported on the nuclear data) within *G. pusillus*, with the level of differentiation (*p*-distances of 10-11%) favoring their species status. First one consists of populations from Eastern Ethiopia; the second includes populations from Kenya and Southern Ethiopia; and the individual from Tanzania represents third lineage (Fig. 2). Within the frame of the current study, we refrain from giving even preliminary names for the identified lineages within *G. pusillus*, because of the absence of genetic data from the type specimens of *G. pusillus* and *G. harwoodi*, apparently being the second species within the subgenus. One can assume that utilizing material from museum collections in further studies can untangle the cryptic diversity within this group.

It is worth noting that such phylogeographic pattern, when populations from southern part of Ethiopia reveal genetic resemblance not with the populations from other parts of the country, but with the conspecifics from neighboring Kenya, is similar to other gerbils (*Gerbilliscus phillipsi* and *G. nigricaudus*, Aghová et al., 2017) as well as other rodents living in the Somali-Masai savanna, e.g. spiny mice *Acomys* (*A. wilsoni* and *A. cahirinus*, Aghová et al., 2019), African grass rats *Arvicanthis* (Bryja et al., 2019), White-bellied rocky mouse *Ochromyscus niveiventris* (Meheretu et al., 2024) and naked mole-rat *Heterocephalus glaber* (Zemlemerova et al., 2021). Taken together, these evidences testify to strong biogeographical affinities of the southern Ethiopia and savannah habitats of the northern Kenya and allow us to consider this micro-region as one of the evolutionary diversity centers within the Horn of Africa.

## Acknowledgements

This study was supported by the Czech Science Foundation, project no. 23-06116S. All fieldwork complied with laws and regulations of Ethiopia, and the sampling was carried out with the permission of the Ethiopian Wildlife Conservation Authority (permission No. EWCA, Ref.No. 31/311/2011). We are grateful Tomáš Mazuch for providing photo and tissue sample of *M. peeli* individual from Somaliland.

## Author contributions

LAL and JB conceived the study and acquired the funding, DSK, AAM, MW, MY, BJ, LAL collected the material in the field; AAM and BP performed the genotyping; EVC performed the chromosomal analysis; AAM analyzed the data; DSK wrote the first draft of the manuscript. All authors contributed to the manuscript and approved its final version.

---

**Fig. S1.**
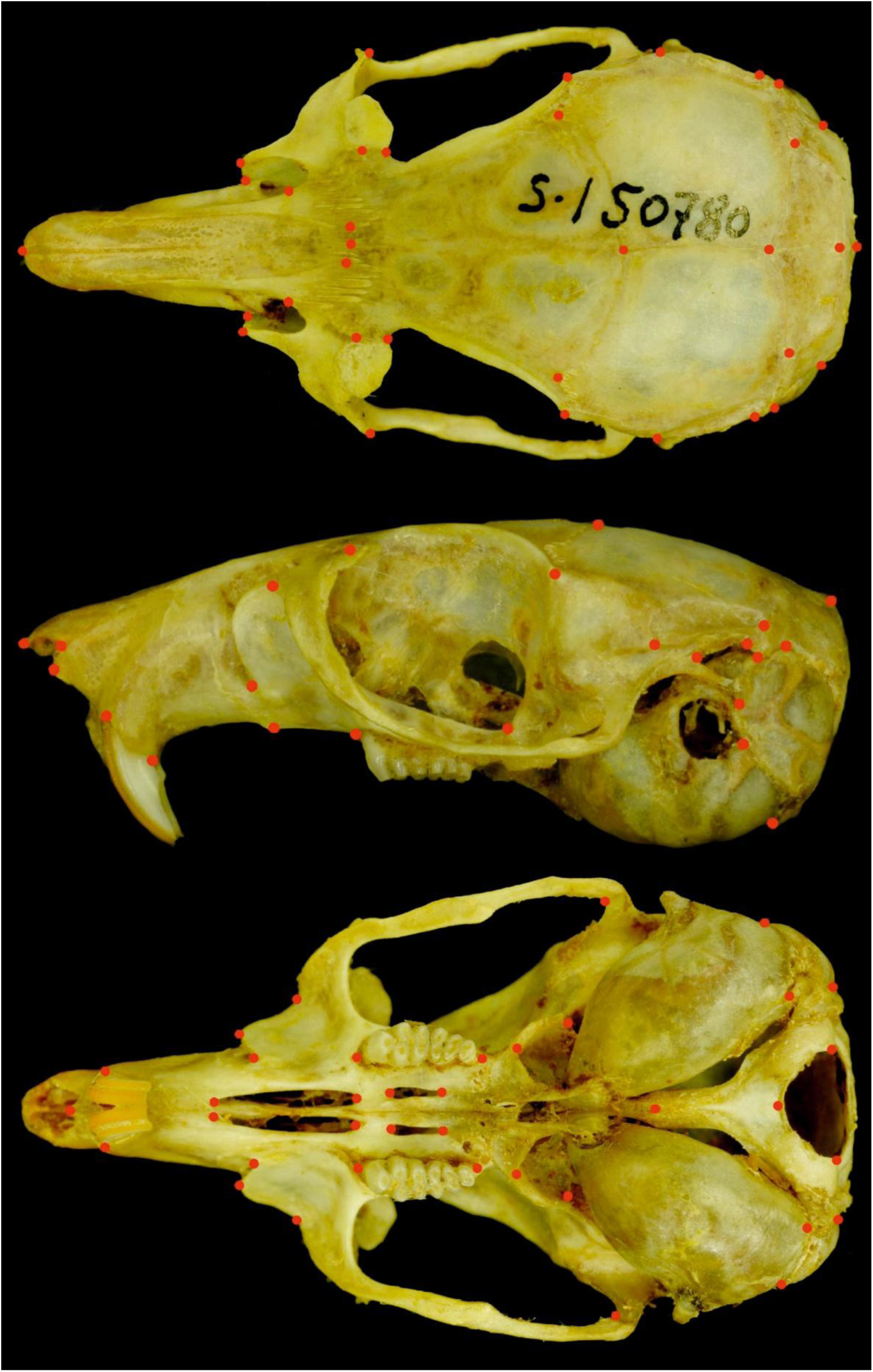
Landmark position scheme

**Fig. S2.**
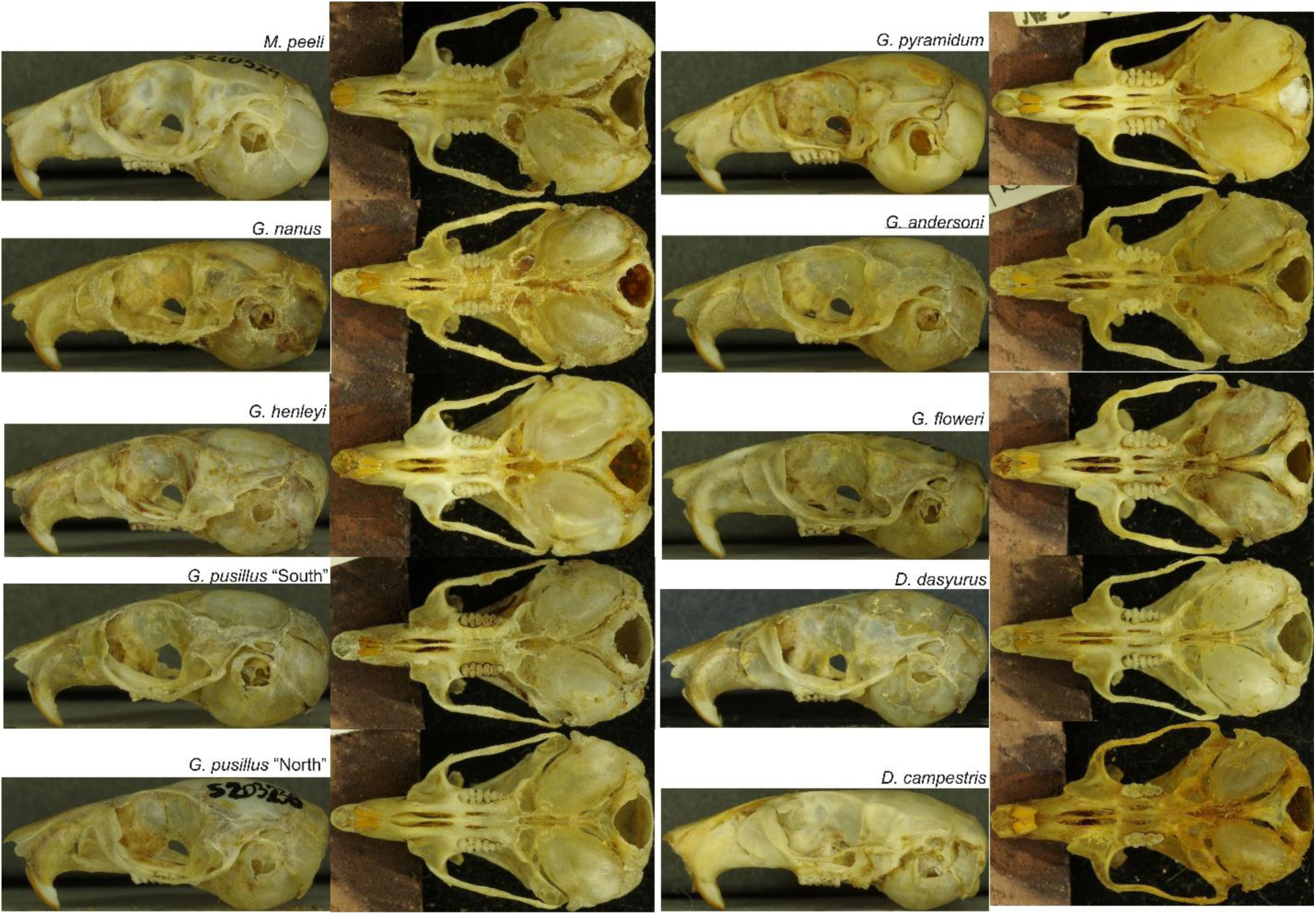
Rescaled images of the dorsal and lateral skull projections for the representatives of the different subgenus within *Gerbillus*: *Microdillus* (*M. peeli*), *Hendecapleura* (*G. nanus*, *G. henleyi*), *Pusillus* (*G. pusillus*), *Gerbillus* (*G. pyramidum*, *G. andersoni*, *G. floweri*), and *Dipodillus* (*D. dasyurus*, *D. campestris*).

**Table S1.** Data on *cytb* sequences used in the study.

| Species | cytb GeneBank ID |
| --- | --- |
| *Ammodillus imbellis* | ON366727 |
| *Brachiones przewalskii* | AB381903 |
| *Desmodilliscus braueri* | AJ851273 |
| *Desmodillus auricularis* | KR089015 |
| *Dipodillus campestris* | KM236116 |
| *Dipodillus campestris* | KF496218 |
| *Dipodillus campestris* | XXX |
| *Dipodillus campestris* | XXX |
| *Dipodillus dasyurus* | OM324328 |
| *Dipodillus dasyurus* | LN606678 |
| *Dipodillus simoni* | GU356579 |
| *Dipodillus simoni* | MH660910 |
| *Gerbillus andersoni* | KF496286 |
| *Gerbillus floweri* | XXX |
| *Gerbillus floweri* | XXX |
| *Gerbillus gerbillus* | KR089023 |
| *Gerbillus gerbillus* | XXX |
| *Gerbillus henleyi* | KF496235 |
| *Gerbillus henleyi* | XXX |
| *Gerbillus henleyi* | XXX |
| *Gerbillus hesperinus* | JN652803 |
| *Gerbillus hoogstrali* | JN021420 |
| *Gerbillus latastei* | GU356550 |
| *Gerbillus nancillus* | KF496264 |
| *Gerbillus nancillus* | KF496258 |
| *Gerbillus nancillus* | KF496241 |
| *Gerbillus nancillus* | KF496279 |
| *Gerbillus nanus* | KR089024 |
| *Gerbillus nanus* | XXX |
| *Gerbillus nigeriae* | AJ430555 |
| *Gerbillus occiduus* | KF496282 |
| *Gerbillus perpallidus* | JN652806 |
| *Gerbillus poecilops* | JQ753064 |
| *Gerbillus pusillus* 'Ethiopian lineage' | XXX |
| *Gerbillus pusillus* 'Ethiopian lineage' | XXX |
| *Gerbillus pusillus* 'Ethiopian lineage' | XXX |
| *Gerbillus pusillus* 'Ethiopian lineage' | XXX |
| *Gerbillus pusillus* 'Ethiopian lineage' | ON366718 |
| *Gerbillus pusillus* 'Ethiopian lineage' | ON366719 |
| *Gerbillus pusillus* 'Ethiopian lineage' | ON366720 |
| *Gerbillus pusillus* 'Ethiopian lineage' | ON366722 |
| *Gerbillus pusillus* 'Ethiopian lineage' | ON366723 |
| *Gerbillus pusillus* 'Ethiopian lineage' | ON366724 |
| *Gerbillus pusillus* 'Ethiopian lineage' | ON366725 |
| *Gerbillus pusillus* 'Ethiopian lineage' | ON366721 |
| *Gerbillus pusillus* 'Ethiopian lineage' | ON366716 |
| *Gerbillus pusillus* 'Ethiopian lineage' | ON366717 |
| *Gerbillus pusillus* 'Ethio-Kenyan lineage' | OL828230 |
| *Gerbillus pusillus* 'Ethio-Kenyan lineage' | OL828231 |
| *Gerbillus pusillus* 'Ethio-Kenyan lineage' | ON366733 |
| *Gerbillus pusillus* 'Tanzanian lineage' | ON366732 |
| *Gerbillus pyramidum* | KF496283 |
| *Gerbillus* sp. | KM236142 |
| *Gerbillus tarabuli* | GU356571 |
| *Meriones tamariscinus* | AB381904 |
| *Microdillus peeli* | XXX |
| *Microdillus peeli* | XXX |
| *Microdillus peeli* | XXX |
| *Microdillus peeli* | XXX |
| *Pachyuromys duprasi* | AJ851274 |
| *Pachyuromys duprasi* | OL828228 |
| *Psammomys obesus* | KR089035 |
| *Psammomys vexillaris* | AY934541 |
| *Rhombomys opimus* | AJ430556 |
| *Sekeetamys calurus* | OL828229 |
| *Sekeetamys calurus* | AJ851276 |

**Table S2.** Data on nuclear markers sequences used in the study.

| Species | IRBP | RAG1 | GHR | ACP5 |
| --- | --- | --- | --- | --- |
| *Ammodillus imbellis* | OL677478 | OL677484 | OL677490 | ON286754 |
| *Ammodillus imbellis* | OL677479 | OL677485 | OL677491 | ON286755 |
| *Desmodillus auricularis* | AM910940 | KC953494 | DQ019048 | KR088977 |
| *Dipodillus campestris* | LN606649 | KR089045 | KR089038 | KR088978 |
| *Dipodillus campestris* | XXX | XXX | XXX | XXX |
| *Dipodillus campestris* | XXX | XXX | XXX | XXX |
| *Dipodillus dasyurus* | LN606652 | KR089046 | KR089039 | KR088979 |
| *Dipodillus simoni* | LN606670 | KR089047 | KR089040 | KR088980 |
| *Gerbillus floweri* | XXX | XXX | XXX | XXX |
| *Gerbillus floweri* | XXX | XXX | XXX | XXX |
| *Gerbillus gerbillus* | EU349846 | DQ023452 | DQ019049 | KR088986 |
| *Gerbillus gerbillus* | XXX | XXX | XXX | XXX |
| *Gerbillus henleyi* | XXX | XXX | XXX | XXX |
| *Gerbillus henleyi* | XXX | XXX | XXX | XXX |
| *Gerbillus nancillus* | LN606658 |  |  | ZZZ |
| *Gerbillus nanus* | LN606660 | KC953501 | KC953262 | KR088987 |
| *Gerbillus nanus* | XXX | XXX | XXX | XXX |
| *Gerbillus nanus* | XXX | XXX | XXX | XXX |
| *Gerbillus pusillus 'Ethiopian lineage'* | XXX | XXX | XXX | XXX |
| *Gerbillus pusillus 'Ethiopian lineage'* | XXX | XXX | XXX | XXX |
| *Gerbillus pusillus 'Ethiopian lineage'* | XXX | XXX | XXX | XXX |
| *Gerbillus pusillus 'Ethiopian lineage'* | XXX | XXX | XXX | XXX |
| *Gerbillus pusillus 'Ethio-Kenyan lineage'* | OL677482 | OL677488 | OL677494 | ON286759 |
| *Gerbillus pusillus 'Ethio-Kenyan lineage'* | XXX | XXX | XXX | XXX |
| *Gerbillus pusillus 'Ethio-Kenyan lineage'* | XXX | XXX | XXX | XXX |
| *Gerbillus pusillus 'Tanzanian lineage'* | XXX | XXX |  | XXX |
| *Meriones meridianus* | OQ507348 | OQ507343 | OQ507353 | OQ507361 |
| *Microdillus peeli* | XXX | XXX | XXX | XXX |
| *Microdillus peeli* | XXX | XXX | XXX | XXX |
| *Microdillus peeli* | XXX | XXX | XXX | XXX |
| *Pachyuromys duprasi* | OL677481 | OL677487 | OL677493 | ON286756 |
| *Psammomys obesus* | FN357290 | MF097927 | MF074924 | KR088996 |
| *Rhombomys opimus* | OQ507345 | OQ507340 | OQ507350 | OQ507359 |
| *Sekeetamys calurus* | OL677481 | OL677486 | OL677492 | ON286757 |
| *Taterillus emini* | KC953461 | DQ023453 | DQ019050 | KR089000 |

**Table S3.** Data on skulls in the morphological analysis.

| Species | ZMMU code | sex | Country | Locality |
| --- | --- | --- | --- | --- |
| *Gerbillus andersoni* | s165632 | m | Israel | Negev desert |
|  | s165634 | f | Israel | Negev desert |
|  | s165637 | m | Israel | Negev desert |
|  | s165639 | m | Israel | Negev desert |
|  | s165642 | f | Israel | Negev desert |
|  | s165643 | m | Israel | Negev desert |
|  | s165645 | m | Israel | Negev desert |
|  | s165646 | f | Israel | Negev desert |
|  | s165647 | m | Israel | Negev desert |
|  | s165648 | m | Israel | Negev desert |
| *Gerbillus campestris* | s150364 | m | Russia | Moscow Zoo |
|  | s153106 | m | Russia | Moscow Zoo |
|  | s153107 | f | Russia | Moscow Zoo |
|  | s156792 | m | Russia | Moscow Zoo |
|  | s156793 | m | Russia | Moscow Zoo |
|  | s157825 | m | Russia | Moscow Zoo |
|  | s158551 | f | Russia | Moscow Zoo |
|  | s161682 | m | Russia | Moscow Zoo |
|  | s175695 |  | Russia | Moscow Zoo |
| *Gerbillus floweri* | s150369 |  | Russia | Moscow Zoo |
|  | s150780 | m | Russia | Moscow Zoo |
|  | s155368 | m | Russia | Moscow Zoo |
|  | s155699 | m | Russia | Moscow Zoo |
|  | s155701 | m | Russia | Moscow Zoo |
|  | s157082 |  | Russia | Moscow Zoo |
|  | s157844 | m | Russia | Moscow Zoo |
|  | s158552 | f | Russia | Moscow Zoo |
|  | s159183 | f | Russia | Moscow Zoo |
| *Gerbillus henleyi* | s165649 | m | Israel | Negev desert |
|  | s165650 | f | Israel | Negev desert |
|  | s165662 | m | Israel | Negev desert |
|  | s165663 | m | Israel | Negev desert |
|  | s165664 | m | Israel | Negev desert |
|  | s165665 | m | Israel | Negev desert |
|  | s165666 | f | Israel | Negev desert |
|  | s165714 |  | Israel | Negev desert |
|  | s165715 | f | Israel | Negev desert |
|  | s179208 | f | Israel | Negev desert |
| *Gerbillus nanus* | s92445 | f | Iran |  |
|  | s169221 | m | Egypt | Aerva |
| *Gerbillus pusillus* | s203219 | m | Ethiopia | Jeldessa |
| 'Ethiopian lineage' | s203231 | m | Ethiopia | Jeldessa |
|  | s203232 | m | Ethiopia | Jeldessa |
|  | s203233 | f | Ethiopia | Jeldessa |
|  | s203235 | f | Ethiopia | Jeldessa |
|  | s203236 | m | Ethiopia | Jeldessa |
|  | s203240 | m | Ethiopia | Jeldessa |
|  | s210411 | f | Ethiopia | Aysha |
|  | s210412 | m | Ethiopia | Aysha |
|  | s210413 | f | Ethiopia | Aysha |
|  | s210414 | m | Ethiopia | Kebri Dahar |
|  | s210415 | m | Ethiopia | Kebri Dahar |
|  | s210416 | f | Ethiopia | Kebri Dahar |
|  | s210417 | m | Ethiopia | Kebri Dahar |
| *Gerbillus pusillus* | s198059 | m | Ethiopia | Megado |
| 'Ethio-Kenyan lineage' | s198060 | m | Ethiopia | Megado |
|  | s198061 | m | Ethiopia | Megado |
|  | s198062 | m | Ethiopia | Megado |
|  | s198063 | f | Ethiopia | Megado |
| *Gerbillus pyramidum* | s8954 | m | Sudan | Port Sudan |
|  | s8956 | m | Sudan | Port Sudan |
|  | s8957 | f | Sudan | Port Sudan |
|  | s8958 | m | Sudan | Port Sudan |
| *Gerbillus dasyurus* | s165620 | m | Israel | Negev desert |
|  | s165621 | f | Israel | Negev desert |
|  | s165622 | f | Israel | Negev desert |
|  | s165623 | f | Israel | Negev desert |
|  | s165625 | f | Israel | Negev desert |
|  | s165627 | f | Israel | Negev desert |
|  | s165675 | f | Israel | Negev desert |
|  | s165676 | m | Israel | Negev desert |
|  | s165713 | f | Israel | Negev desert |
|  | s179194 | m | Israel | Negev desert |
| *Microdillus peeli* | s210522 | f | Ethiopia | Aysha |
|  | s210523 | f | Ethiopia | Aysha |
|  | s210524 | m | Ethiopia | Aysha |

**Table S4.**
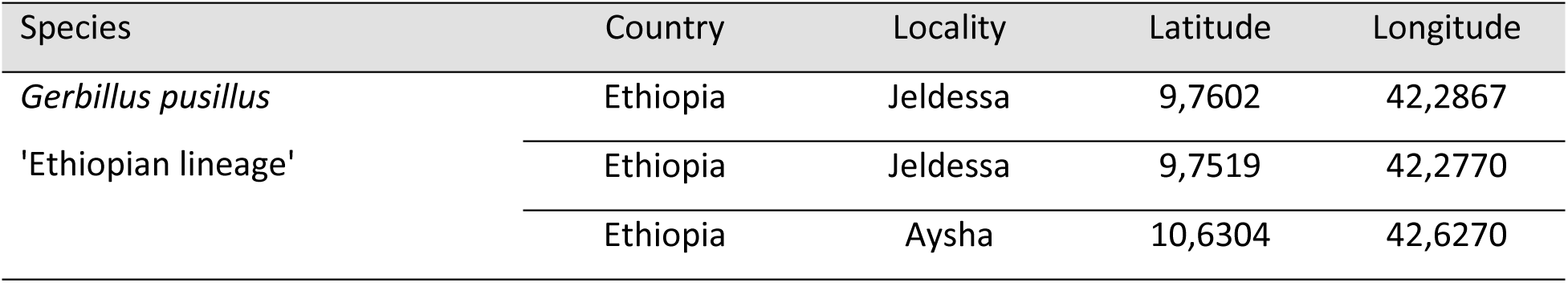
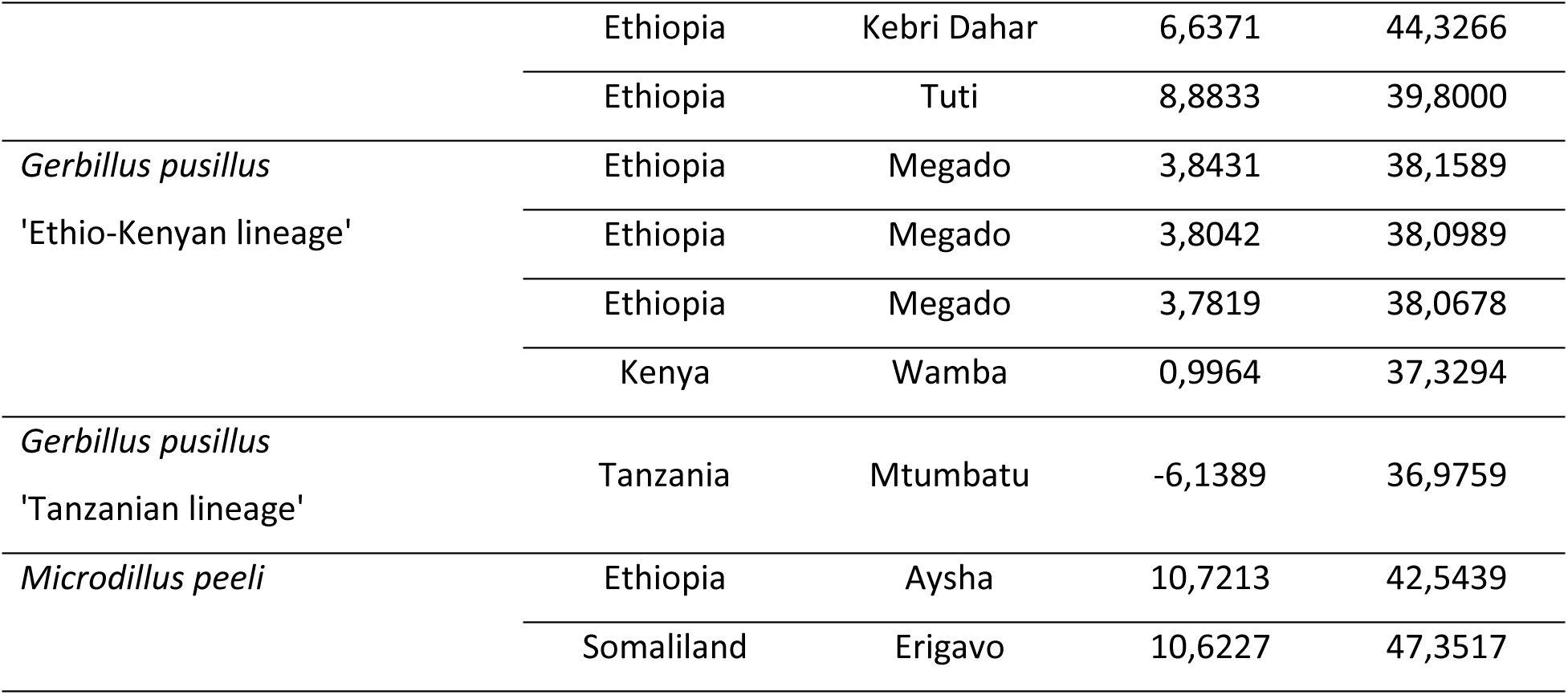
Individual samples collection sites.

## Notes

### Competing Interest Statement

The authors have declared no competing interest.

## References

Adkins, R. M., Gelke, E. L., Rowe, D., & Honeycutt, R. L. (2001). Molecular phylogeny and divergence time estimates for major rodent groups: evidence from multiple genes. Molecular Biology and Evolution, 18(5), 777–791.

Aghova, T., Palupčikova, K., Šumbera, R., Frynta, D., Lavrenchenko, L. A., Meheretu, Y., Sadlova, J., Votypka, J., Mbau, J. S., Modry, D., & Bryja, J. (2019). Multiple radiations of spiny mice (Rodentia: *Acomys*) in dry open habitats of afro-Arabia: Evidence from a multi-locus phylogeny. BMC Evolutionary Biology, 19(1), 1–22.

Aghova, T., Šumbera, R., Pialek, L., Mikula, O., McDonough, M. M., Lavrenchenko, L. A., Meheretu, Y., Mbau, J. S., & Bryja, J. (2017). Multilocus phylogeny of east African gerbils (Rodentia, *Gerbilliscus*) illuminates the history of the Somali-Masai savannah. Journal of Biogeography, 44(10), 2295–2307.

Alhajeri, B. H. (2018). Craniomandibular variation in the taxonomically problematic gerbil genus Gerbillus (Gerbillinae, Rodentia): assessing the influence of climate, geography, phylogeny, and size. Journal of Mammalian Evolution, 25(2), 261–276.

Bouckaert, R., Vaughan, T. G., Barido-Sottani, J., Duchêne, S., Fourment, M., Gavryushkina, A., & Drummond, A. J. (2019). BEAST 2.5: An advanced software platform for Bayesian evolutionary analysis. PLoS computational biology, 15(4), e1006650.

Bryja, J., Colangelo, P., Lavrenchenko, L. A., Meheretu, Y., Šumbera, R., Bryjová, A., Verheyen, E., Leirs, H., & Castiglia, R. (2019). Diversity and evolution of African grass rats (Muridae: *Arvicanthis*)—from radiation in East Africa to repeated colonization of northwestern and southeastern savannas. Journal of Zoological Systematics and Evolutionary Research, 57(4), 970–988.

Bryja, J., Meheretu, Y., Boratyński, Z., Zeynu, A., Denys, C., Mulualem, G., Welegerima, K., Bryjova, A., Kasso, M., Kostin, D. S., Martynov, A. A., & Lavrenchenko, L. A. (2022). Rodents of the Afar triangle (Ethiopia): Geographical isolation causes high level of endemism. Biodiversity and Conservation, 31, 629–650.

Capanna, E., & Merani, M. S. (1981). Karyotypes of somalian rodent populations: 2. The chromosomes of *Gerbillus dunni* (Phomas, 1904), *Gerbillus pusillus* (Peters, 1878) and *Ammodillus imbellis* (de Winton, 1898) (cricetidae gerbillinae): Pubblicazioni del centro di studio per la faunistica ed ecologia tropicali del cnr: CCII. Monitore Zoologico Italiano, 14(1), 227–240.

Chevret, P., & Dobigny, G. (2005). Systematics and evolution of the subfamily Gerbillinae (Mammalia, Rodentia, Muridae). Molecular phylogenetics and evolution, 35(3), 674–688.

Cui, Y., Cheng, J., Wen, Z., Feijó, A., Xia, L., Ge, D., Artige, E., Granjon, L., & Yang, Q. (2025). Evolutionary factors and habitat filtering affect the pattern of Gerbillinae diversity. Current zoology, 71(1), 65–78.

DeBry, R. W., & Seshadri, S. (2001). Nuclear intron sequences for phylogenetics of closely related mammals: an example using the phylogeny of *Mus*. Journal of Mammalogy, 82(2), 280–288.

de Winton, W. E. (1898). XXXIX.—On a small collection of mammals made by Mr. CVA Peel in Somaliland. Journal of Natural History, 1(3), 247–251.

Douglas, J., Jiménez-Silva, C. L., & Bouckaert, R. (2022). StarBeast3: adaptive parallelized Bayesian inference under the multispecies coalescent. Systematic Biology, 71(4), 901–916.

Ford, C. E., & Hamerton, J. L. (1956). A colchicine, hypotonic citrate, squash sequence for mammalian chromosomes. Stain Technology, 31, 247–251.

Granjon, L. 2013. Genus *Gerbillus*; pp 295–297 in Happold, D. C. D. (ed.) 2013. Mammals of Africa: Volume III. Bloomsbury Publishing, London.

Hall, T. A. (1999, January). BioEdit: a user-friendly biological sequence alignment editor and analysis program for Windows 95/98/NT. In Nucleic acids symposium series (Vol. 41, No. 41, pp. 95–98).

Hammer, Ø., & Harper, D. A. (2001). Past: paleontological statistics software package for educaton and data anlysis. Palaeontologia electronica, 4(1), 1.

Kalyaanamoorthy, S., Minh, B. Q., Wong, T. K., Von Haeseler, A., & Jermiin, L. S. (2017). ModelFinder: Fast model selection for accurate phylogenetic estimates. Nature Methods, 14(6), 587–589.

Kostin, D. S., Martynov, A. A., Lebedev, V. S., Zemlemerova, E. D., Gromov, A. R., & Lavrenchenko, L. A. (2022). Position of the ammodile and the origin of Gerbillinae (Rodentia): Out of the Horn of Africa?. Zoologica Scripta, 51(5), 522–532.

Lanfear, R., Frandsen, P. B., Wright, A. M., Senfeld, T., & Calcott, B. (2017). PartitionFinder 2: new methods for selecting partitioned models of evolution for molecular and morphological phylogenetic analyses. Molecular biology and evolution, 34(3), 772–773.

Lecompte, É., Granjon, L., Peterhans, J. K., & Denys, C. (2002). Cytochrome *b*-based phylogeny of the *Praomys* group (Rodentia, Murinae): a new African radiation?. Comptes rendus. Biologies, 325(7), 827–840.

Mammal Diversity Database. (2025). Mammal Diversity Database (v2.2) [Data set]. Zenodo. 10.5281/zenodo.15659662

Manthi, F. K. (2007). A preliminary review of the rodent fauna from Lemudong’o, southwestern Kenya, and its implication to the late Miocene paleoenvironments. Kirtlandia, 56, 92–105.

Meheretu, Y., Mikula, O., Frynta, D., Frýdlová, P., Mulualem, G., Lavrenchenko, L. A., Kostin, D. S., Abdirahman, H. E., Šumbera, R., & Bryja, J. (2024). Phylogeny, biogeography, and integrative taxonomic revision of the Afro-Arabian rodent genus *Ochromyscus* (Muridae: Murinae: Praomyini). Zoological Journal of the Linnean Society, 202(1), zlad158.

Ndiaye, A., Ba, K., Aniskin, V., Benazzou, T., Chevret, P., Konečný, A., Sembene, M., Tatard, C., Kergoat, G. J., & Granjon, L. (2012). Evolutionary systematics and biogeography of endemic gerbils (Rodentia, Muridae) from Morocco: an integrative approach. Zoologica Scripta, 41(1), 11–28.

Ndiaye, A., Chevret, P., Dobigny, G., & Granjon, L. (2016). Evolutionary systematics and biogeography of the arid habitat-adapted rodent genus *Gerbillus* (Rodentia, Muridae): A mostly Plio-Pleistocene African history. Journal of Zoological Systematics and Evolutionary Research, 54(4), 299–317.

Nguyen, L. T., Schmidt, H. A., Von Haeseler, A., & Minh, B. Q. (2015). IQ-TREE: a fast and effective stochastic algorithm for estimating maximum-likelihood phylogenies. Molecular biology and evolution, 32(1), 268–274.

Pavlinov, I. Y. (2008). A review of phylogeny and classification of Gerbillinae (Mammalia: Rodentia) (p. 68). Moscow University Publishing.

Pavlinov, I. Y., Dubrovsky, Y. A., Rossolimo, O. L., & Potapova, E. G. (1990). Gerbillides of the world (p. 368). Nauka Publ.

Piwczyński, M., Granjon, L., Trzeciak, P., Brito, J. C., Popa, M. O., Dinka, M. D., Johnston, N. P., & Boratyński, Z. (2023). Unraveling phylogenetic relationships and species boundaries in the arid adapted *Gerbillus* rodents (Muridae: Gerbillinae) by RAD-seq data. Molecular phylogenetics and evolution, 189, 107913.

Rambaut, A., Suchard, M. A., Xie, D. & Drummond, A. J. (2014). Tracer v1. 6. http://beast.bio.ed.ac.uk/Tracer

Roche, J. (1975). A propos des petites gerbilles a soles plantaires nues (sous-genre *Hendecapleura*) de l’est africain: pubblicazioni del centro di studio per la faunistica ed ecologia tropicali del cnr: XCIX. Monitore Zoologico Italiano. Supplemento, 6(1), 263–268.

Rohlf, F.J. (2007). tpsRelw, relative warps analysis, version 1.45. Department of Ecology and Evolution, State University of New York, Stony Brook.

Rohlf, F.J. (2008). tpsDig, digitize landmarks and outlines, version 2.12. Department of Ecology and Evolution, State University of New York, Stony Brook.

Rohlf, F.J. and Slice, D. (1990). Extensions of the Procrustes method for the optimal superimposition of landmarks. Syst. Zool. 39: 40–59.

Ronquist, F., Teslenko, M., Van Der Mark, P., Ayres, D.L., Darling, A., Höhna, S., Larget, B., Liu, L., Suchard, M.A., and Huelsenbeck, J.P. (2012). MrBayes 3.2: efficient Bayesian phylogenetic inference and model choice across a large model space. Syst. Biol. 61: 539–542.

Stanhope, M. J., Czelusniak, J., Si, J. S., Nickerson, J., & Goodman, M. (1992). A molecular perspective on mammalian evolution from the gene encoding interphotoreceptor retinoid binding protein, with convincing evidence for bat monophyly. Molecular Phylogenetics and Evolution, 1(2), 148–160.

Zemlemerova, E. D., Kostin, D. S., Lebedev, V. S., Martynov, A. A., Gromov, A. R., Alexandrov, D. Y., & Lavrenchenko, L. A. (2021). Genetic diversity of the naked mole-rat (*Heterocephalus glaber*). Journal of Zoological Systematics and Evolutionary Research, 59(1), 323–340.

